# dNEMO: a tool for quantification of mRNA and punctate structures in time-lapse images of single cells

**DOI:** 10.1101/855213

**Authors:** Gabriel J. Kowalczyk, J. Agustin Cruz, Yue Guo, Qiuhong Zhang, Natalie Sauerwald, Robin E. C. Lee

## Abstract

Many biological processes are regulated by single molecules and molecular assemblies within cells that are visible by microscopy as punctate features, often diffraction limited. Here we present detecting-NEMO (dNEMO), a computational tool optimized for accurate and rapid measurement of fluorescent puncta in fixed-cell and time-lapse images. The spot detection algorithm uses the *à trous* wavelet transform, a computationally inexpensive method that is robust to imaging noise. By combining automated with manual spot curation in the user interface, fluorescent puncta can be carefully selected and measured against their local background to extract high quality single-cell data. Integrated into the workflow are segmentation and spot-inspection tools that enable almost real-time interaction with images without time consuming pre-processing steps. Although the software is agnostic to the type of puncta imaged, we demonstrate dNEMO using smFISH to measure transcript numbers in single cells in addition to the transient formation of IKK/NEMO puncta from time-lapse images of cells exposed to inflammatory stimuli.

## INTRODUCTION

Quantitative imaging of single cells enables measurement with subcellular resolution of dynamic biological processes that regulate critical cellular behaviors. Processes of the central dogma such as active transcription, single mRNA transcripts, sites of active protein translation, and other regulatory multi-protein assemblies can be observed by fluorescence microscopy as punctate structures within the cell (1-6). Spatiotemporal dynamics of signaling proteins, quantified in single cells by live-cell imaging, have revealed mechanistic insights into signal-response relationships in signal transduction networks in addition to sources of cell-to-cell variability (7-9). However, most live-cell imaging approaches use fluorescent biosensors and fusion proteins that report within large subcellular compartments, such as the cytoplasm or nucleus, and quantification of punctate structures is often limited to fixed-cell and low-throughput applications. Accurate detection and quantification of biological puncta is necessary to examine their roles in regulating cellular behaviors, and computational analysis is often the rate-limiting step of experimental pipelines.

The nuclear factor (NF)-κB signal transduction pathway is a master regulator of inflammatory responses to injury and infection (10). Following activation of the pathway by inflammatory cytokines, such as interleukin-1 (IL-1) or tumor necrosis factor (TNF) among others, a series of intracellular signaling events transduce the NF-κB signal. Upstream kinase activation by the NF-κB essential modulator protein (NEMO, also known as IKKγ) is a necessary step in regulation of the classical NF-κB signaling cascade (11). Following cytokine stimulation, NEMO is recruited to polyubiquitin scaffolds associated with cytokine-ligated receptor complexes where NEMO-interacting IκB kinases (IKKs) are activated through induced proximity with other signaling mediators (12-16). In cells that express EGFP fused to NEMO exposed to inflammatory cytokines, EGFP-NEMO transiently localizes to punctate fluorescent structures near the plasma membrane (16, 17). The number and timescales of EGFP-NEMO-enriched puncta reveal quantitative properties of receptor-associated protein complexes that transmit information from the inflammatory milieu into the NF-κB transcriptional system.

Here we present detecting NEMO (dNEMO), a free application that uses wavelet-based spot detection and supervised segmentation to detect and measure properties of fluorescent puncta in fixed-cell and time-lapse images. We show that the wavelet-based approach is significantly faster than traditional Gaussian fitting methods and allows for almost real-time interaction with single cells in quantitative imaging data. Intuitive tools for cell segmentation, spot inspection, and background correction, in addition to manual and automated selection of puncta based on quantifiable features (e.g. size, location, fluorescence) ensure that single cell data are of the highest quality. Results are formatted for easy coordination with other software packages, such as single-particle tracking applications and other analyses of structural dynamics (18). Using smFISH and live-cell data for EGFP-NEMO as demonstrations, we show that dNEMO is a versatile workspace for rapid, precise, and robust measurement of fluorescent puncta in digital images.

## RESULTS

### dNEMO identifies near diffraction-limited fluorescent puncta in 2D and 3D images

Wavelet-based approaches are used in image analysis for de-noising, compression, and feature extraction with low computational cost (19, 20). In wavelet-based feature extraction applications, the source image is decomposed into wavelet maps, a series of images where contrast is enhanced for particular spatial features. Since the wavelet transform sequentially applies a different convolution matrix at successive levels of the algorithm, the size and qualities of spatial features that are enhanced in each wavelet map can be modulated.

The *à trous* wavelet transform accurately detects and localizes isotropic diffraction-limited spots such as fluorescent mRNA puncta in single molecule FISH images (21, 22). As the wavelet map transform level increases, zeros are progressively inserted into the convolution matrix (Figure 1A, see also Supplementary Figure 1). Comparing experimental images of diffraction-limited spots at the first level of the wavelet algorithm (L1 wavelet map), noise and the smallest puncta in the source image were enhanced (Figure 1B, 2^nd^ column). Consistent with previous findings (22, 23), the L2 wavelet map (Figure 1B, 3^rd^ column; see also Movie S1) enhanced contrast for puncta at or near the diffraction limit. At higher levels, larger puncta were more resolved (L3 wavelet map; Figure 1B, 4^th^ column) at the expense of reduced clarity for smaller puncta. Although users of dNEMO can select a wavelet map appropriate for their application, the L2 wavelet map was used for all subsequent experiments to detect small molecular assemblies.

**Figure 1:**
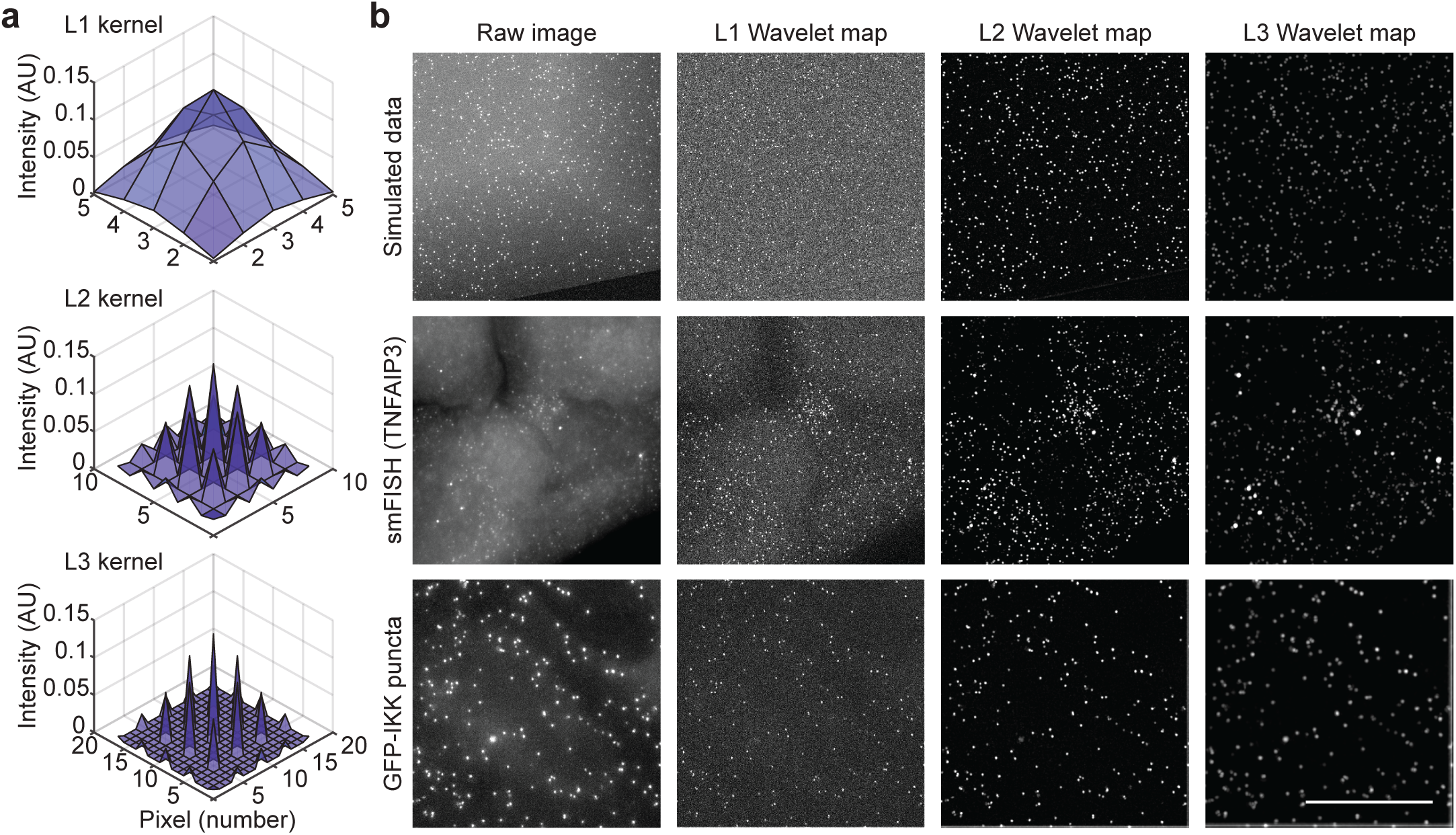
The *à trous* wavelet transform on simulated and experimental images. **(a)** 3D representations of the convolution matrix (kernel) for levels 1 through 3 of the wavelet transform. **(b)** Images for simulated data (top), smFISH-labeled *NFKBIA* transcripts (middle), or GFP-NEMO (bottom), along with the associated L1, L2, and L3 wavelet maps. The L2 wavelet map enhances contrast for diffraction-limited puncta in fluorescence microscopy images. Scale bar 25 microns.

To identify fluorescent puncta near the diffraction limit, the L2 wavelet map was segmented using thresholding and watershed algorithms in dNEMO. Briefly, the background is removed from the L2 wavelet map by setting pixel values below a user-defined threshold to zero and multiplying other pixel values by −1 to generate an inverted wavelet map. A watershed is then applied to define local basins in the inverted wavelet map. The dNEMO interface updates the source image in real-time to assist users with a first-pass visual estimation for puncta identified under the chosen threshold value (Figure 2, top panel). The spatial coordinates for each basin and the surrounding path are used to define the centroid and perimeter respectively for each of the fluorescent punctum. Puncta are then evaluated to prevent over-segmentation, where a single punctum with a noisy spatial distribution of fluorescence is erroneously segmented by the watershed into two or more puncta. To resolve over-segmentation, the paths that connect all pairs of centroids separated by less than 10 pixels are scanned for a minimum intensity value. If a minimum is not found, the puncta are consolidated, and properties of the merged punctum are recalculated. Similarly, centroids falling within 3 pixels of each other are combined because this is within the resolution limit of our optical imaging system.

**Figure 2:**
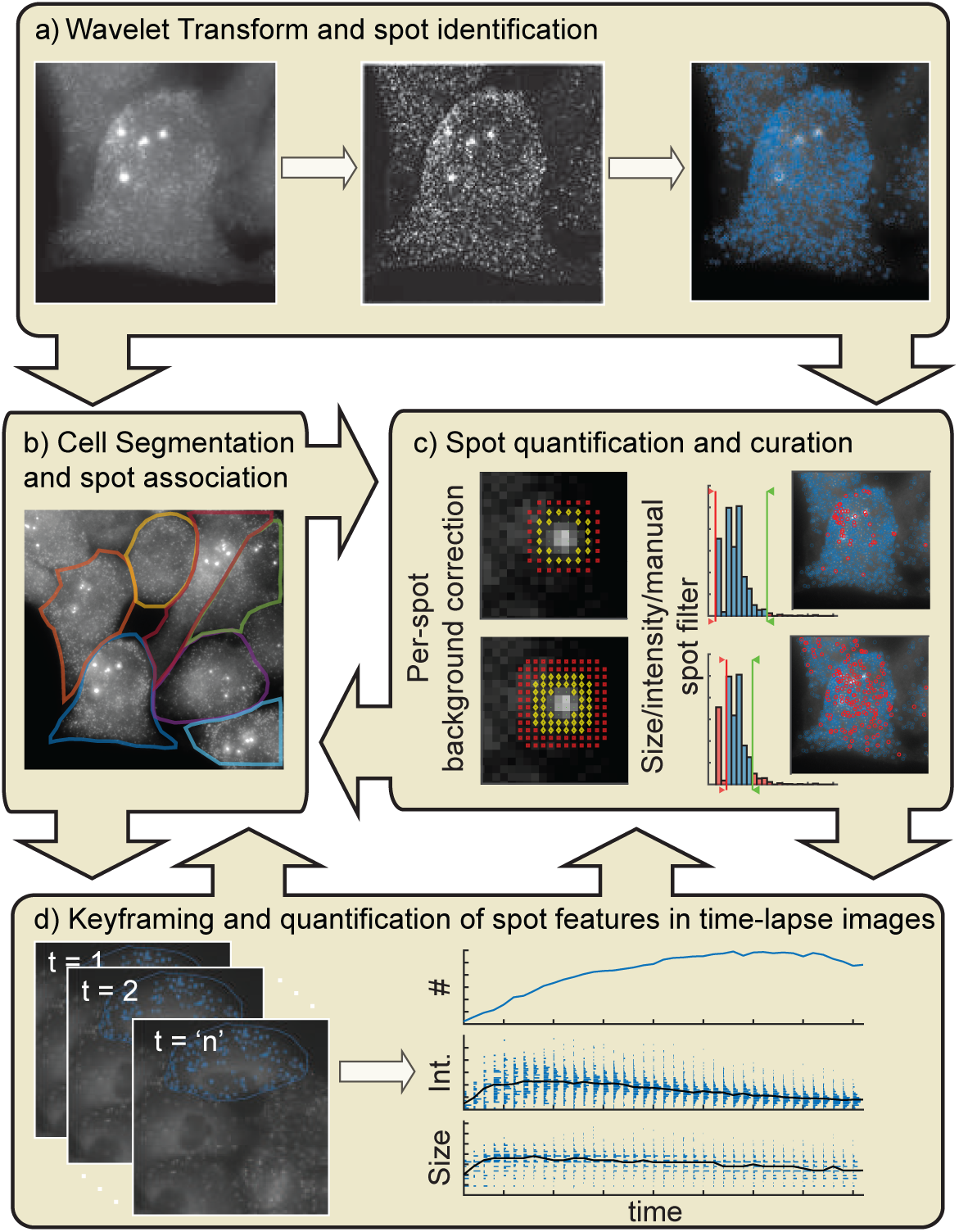
Overview of the dNEMO workflow. **(a)** In the first operation performed by the application, the image undergoes the *à trous* wavelet transform, producing a wavelet map, which is subsequently segmented using by watershed to identify puncta. **(b)** Individual cells are separately identified through an interactive manual segmentation tool operated by the user (middle left). **(c)** Once identified by the wavelet transform, identified puncta can be curated based on features like intensity and size. Puncta intensities are corrected using local background pixels for each individual punctum. **(d)** Settings used to define valid puncta for a single image are propagated over sets of time-lapse images creating a keyframe. Combined with previous segmentation of individual cells, punctum features are quantified over time and associated to single cells.

For analysis of puncta in 3D images, wavelet maps are produced for each 2D slice of the image stack. Centroids identified in each slice are referenced against centroids in adjacent slices and puncta are merged between slices of the image stack if the X-Y coordinates of their centroids fall within a Euclidean distance of 2 pixels. Using this approach in parallel with fluorescence information for the same punctum in adjacent image slices, an axial component of each centroid can be calculated and overlapping spots can be resolved (Supplemental Figure 2). Fluorescence properties for each 3D punctum can be either aggregated across image slices or measured at the image slice that corresponds to the axial centroid. Once user-defined settings are established, dNEMO is typically run in batch processing mode so that the methods for puncta detection are identical across all images in an experiment.

### Local background correction and cell segmentation for accurate quantification of puncta in single cells

Slow-varying background from non-specific dye accumulation and free fluorescent proteins that are not part of molecular assemblies, among other sources, will contribute to the measured intensity of a punctum. To correct for these effects, dNEMO collects local background pixel information for each punctum in the source image. Puncta identified in the inverted wavelet map are dilated (24) to define annular rings around each punctum with user-defined offset and width (Figure 2, middle right panel and Supplemental Figure 3; see also Methods). A distribution of pixel intensities in the source image is measured within each annulus to establish the mean fluorescence intensity of the local background. For pixels identified in puncta, fluorescence intensity values are measured in the source image and the local background is subtracted. To ensure accurate estimation of the local background, annuli pixels are not collected in the vicinity of other puncta (Supplemental Figure 3B). Instead of using procedural generation of annuli for each punctum, the method in dNEMO operates directly on the wavelet map and consequently background correction is rapid.

To associate and compare puncta between single cells, dNEMO contains an interactive polygon tool for manual cell segmentation (Figure 2, middle left panel). All puncta contained within the polygon are associated and puncta features, such as their number and distributions of fluorescent intensities among others, can be collated for each single cell (Figure 2, middle left panel). As a demonstration, we used dNEMO to detect single molecules of mRNA from smFISH images of TNF-induced expression of the *NFKBIA* gene (25). Although there was significant variability in transcript numbers when compared between single cells, the size and fluorescence intensity distributions of puncta were similar (Figure 3).

**Figure 3:**
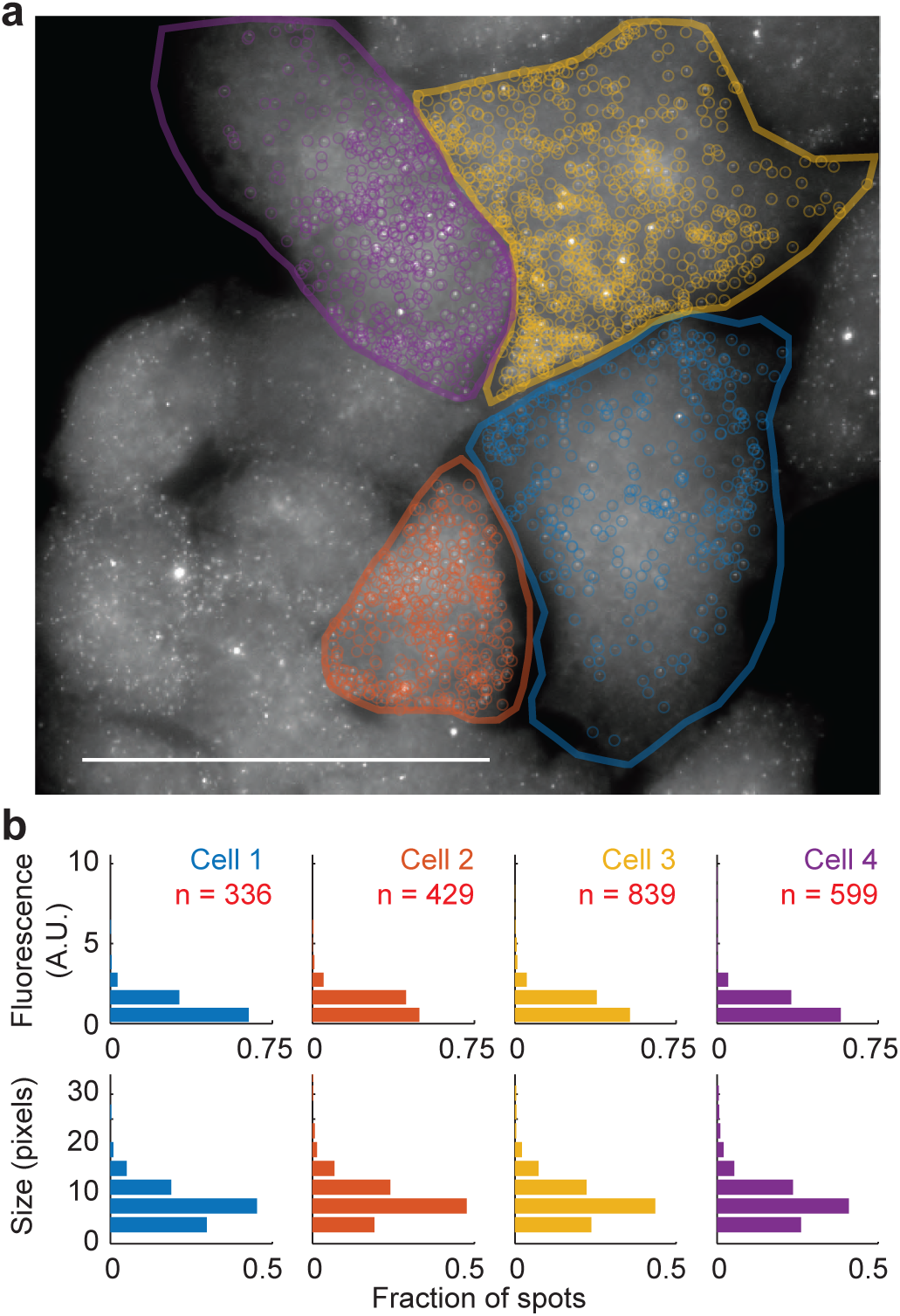
Identification of smFISH transcripts in fixed-cell images. **(a)** *NFKBIA* transcripts in HeLa cells labeled by smFISH are identified using dNEMO and associated to single cells. Scale bar 50 microns. **(b)** Distribution of puncta identified in individual cells reported by intensity (top) or size in number of pixels (bottom). Fluorescence per spot was corrected using the local background about each spot (see Supplemental Figure 3).

In the final analysis for 2D and 3D images, fluorescent and spatial properties are measured for each background-corrected punctum. User-defined bounds for the size and intensity of puncta can be used to filter puncta within a chosen range of fluorescence and size. In the user interface, filtered puncta are indicated with a red circle (Figure 2, middle right panel) which can also be hidden for visual clarity. The interface also provides tools for manual exclusion of erroneously identified puncta, and a spot-inspector tool to examine features of puncta in closer detail. Cell segmentation is independent from the detection and curation of puncta, and features are updated *post hoc* for each segmented cell if user-defined settings for spot detection are changed.

### Spot detection in dNEMO is rapid and accurate

In comparison with spot-fitting methods, such as 3D Gaussian or maximum likelihood estimation (26), the wavelet-based approach in dNEMO does not require iterative estimation of parameters or image pre-processing steps. To compare against our application, we selected the software package FISH-quant (27) primarily because it implements a 3D Gaussian fitting method for detection of transcripts and it’s actively used in the research community. For example, we had previously used FISH-quant to detect single mRNA molecules in the context of stochastic transcription events (28).

Although the localization accuracy and runtime of the *à trous* and Gaussian fitting methods have been compared at the level of the algorithm (22), we set out to make a practical comparison of applications using simulated and fixed-cell data. To test both applications, we generated noisy experimental images using a theoretical point spread function (PSF) to simulate diffraction-limited fluorescent puncta in a slow-varying background with varying amounts of Gaussian noise (Top row, Figure 1b; Supplemental Figure 4a; see also Methods). We compared localization accuracy for dNEMO and FISH-quant using the same sets of simulated images. With ground truth information for the position and number of puncta in simulated data, we found that both applications have almost negligible localization error in low noise, but error rates for dNEMO remained lower for images with greater noise (Supplemental Figure 4b). Wall-clock time for single core performance using whole-image smFISH data showed that dNEMO is over 15-fold faster than FISH-quant on a modern laptop computer (Supplemental Figure 4c), with an estimated 80% elapsed execution time dedicated to the over-segmentation algorithm.

We finally compared accuracy of the background-corrected intensity for detected puncta by comparing intensity values for a raw image with a deconvolved image. Deconvolution is a computationally expensive pre-processing approach that redistributes out-of-focus light in a 3D image, thereby increasing the effective image resolution and precision of intensity measurements. R^2^ values demonstrated that background corrected intensity values for raw and deconvolved images (Supplemental Figure 4d) are almost equivalent, and significantly greater than uncorrected images (R^2^ values of 90 and 58 respectively; Supplemental Figure 4e). We therefore conclude that image pre-processing steps such as deconvolution do not necessarily increase the relative accuracy of dNEMO intensity measurements in arbitrary units. Taken together, when images are collected in the linear regime of an imaging sensor such as the sCMOS used here, spot quantification in dNEMO is rapid, accurate, and robust to noise in imaging data.

### Keyframing to detect time-varying features of puncta in time-lapse images

Keyframing is a process in animation that denotes the start and end frames in a time series where parameter values change. dNEMO uses a keyframing approach for users to make changes for any user-defined parameter and to track single cells in time-lapse experiments. For example, a user may define a region of the time-lapse where the wavelet map threshold or the user-defined bounds for acceptable puncta are modified to account for effects of photobleaching or other systematic artifacts. A more common use for keyframing in dNEMO is to adjust the segmentation polygon to account for morphology changes and cell movement over the time-lapse image (Figure 2, bottom panel).

To demonstrate keyframing we analyzed a time-lapse image of CRISPR/Cas9-modified U2OS cells that express EGFP-tagged NEMO from its endogenous gene locus in response to IL-1(16). Formation of NEMO puncta in single cells were tracked by making keyframe adjustments to cell segmentation polygons (Figure 4A; see also Movie S2). Fluorescent properties of NEMO puncta were followed over the time-lapse to monitor time-courses for adaptive changes in NEMO puncta numbers in addition to distributions for fluorescent properties for NEMO puncta over time in each cell (Figure 4B). By selecting appropriate parameters, such as the wavelet map level and limiting boundaries for puncta intensity or size, punctate structures can be accurately measured and curated in digital images to produce high-quality single-cell datasets.

**Figure 4:**
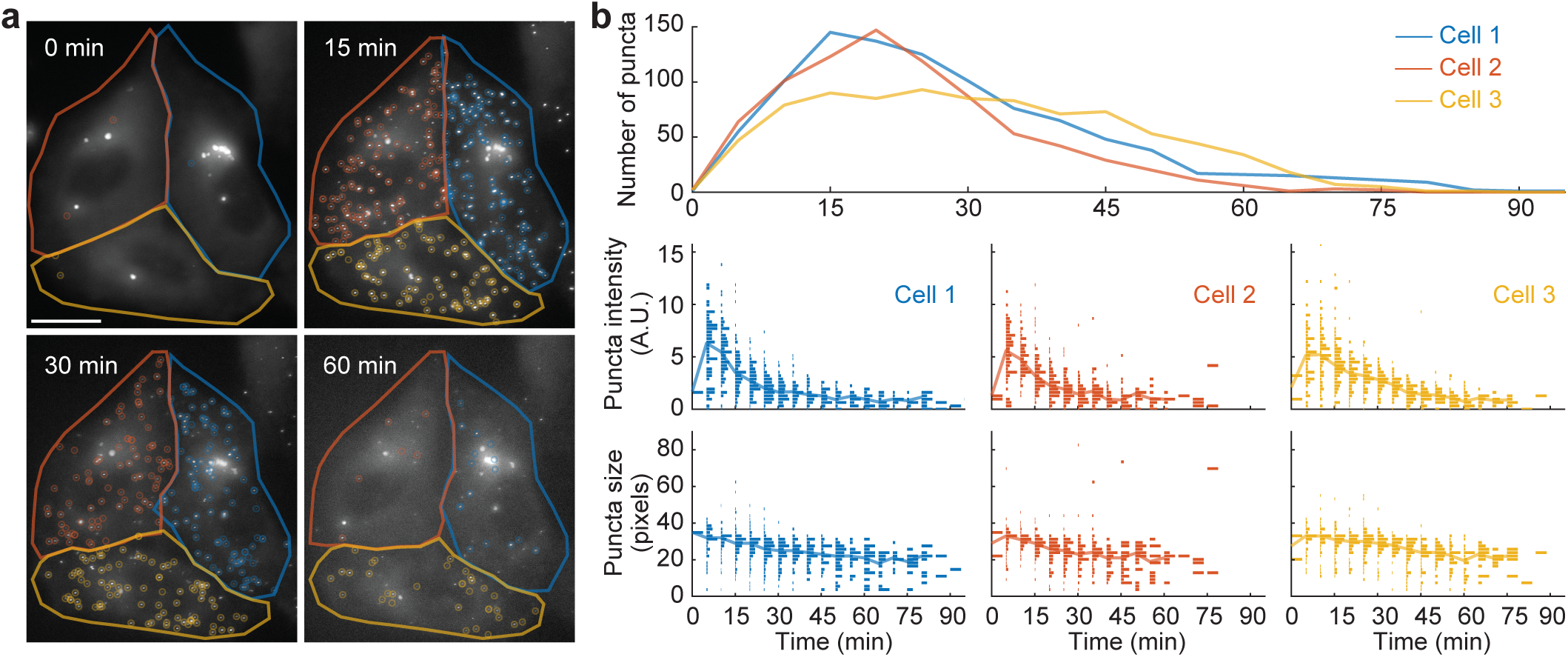
Quantification of EGFP- NEMO in live-cell time-lapse images. **(a)** Live-cell time-lapse images of U2OS cells expressing EGFP-NEMO from its endogenous locus exposed to 100 ng/mL IL-1. NEMO transiently localized to punctate structures and were identified with dNEMO and associated to individual cells. Scale bar 20 microns. **(b)** Analysis of puncta identified within single cells over time. The number of puncta (top), the distribution of puncta intensities (middle), and the distribution of puncta sizes (bottom) are shown per cell over time.

## DISCUSSION

In this work we have shown that dNEMO is an effective tool for quantification of fluorescent puncta in fixed-cell and time-lapse images. The *à trous* wavelet algorithm progressively removes high frequency noise within fluorescent images and can be used to enhance puncta near the diffraction limit and larger. When compared with established methods, dNEMO performs with comparable or better localization accuracy, depending on the amount of noise in the source image, and is significantly faster. Although dNEMO is suitable for detection of relatively bright structures in epifluorescence images, model fitting may still be the preferred method for detection of structures with lower signal-to-noise (29, 30). Keyframing in dNEMO provides an effective interface to curate single cell data and correct for systematic effects in imaging data. We demonstrate dNEMO using fixed-cell smFISH and live-cell enrichment of EGFP-NEMO to puncta and expect dNEMO will also excel at quantifying other fluorescence reporters, including components of the central dogma, protein assemblies, and bright vesicular structures.

Updates to dNEMO are expected to further reduce its runtime and enhance its capabilities. One notable limitation of the current dNEMO implementation is the disproportionate amount of overhead dedicated to the over-segmentation algorithm. We expect that these can be mitigated through updates for parallelization of the over-segmentation process or by modifications to the watershed algorithm that reduce the computational expense while maintaining accuracy. Beyond runtime improvements, one of the largest bottlenecks is the manual segmentation of cells. We are actively considering experimental methods for labeling and incorporating an automated cell segmentation approach into dNEMO, either directly or through a plug-in system where users can choose their own cell segmentation method. Finally, we are also streamlining dNEMO data structures for compatibility with existing single-particle tracking packages (31), so that time-varying properties of single puncta can be tracked and associated with single-cell responses.

Tools dedicated to the processing of biological images have enabled many studies of single cell variability and dynamics, and contributed to the discovery of emergent cellular properties. dNEMO fills a gap in the scientific community by providing a simple workspace for users to interact with biological puncta in fluorescence microscopy images that are central to fundamental cellular processes. The software is controlled with a MATLAB user interface or as a stand-alone executable, and is available as Supplementary Software or at https://github.com/recleelab along with a user manual and test data used to generate the figures in this article.

## METHODS

### The dNEMO user interface

The user interface is a MATLAB-based application which provides several means of interaction with single-channel images and movies. Users load a given image or movie into the application. The two overarching processes of the application, cellular segmentation and puncta identification, are independent of one another and do not rely on each other to properly function. Puncta are identified by creating a keyframe to perform the wavelet transform over the frames of a time-lapse image. Further curation is handled using an interactive distribution of identified puncta within the current image and imposing limits based on spot size or intensity among other measured properties. Settings like the wavelet map level and the minimum number of consecutive slices of a 3D image a spot must appear in are user-defined parameters. Once the user settings are established for an experiment, the same settings are batch processed over all indicated images. A cell segmentation button in the interface initiates a process where the user defines a cell perimeter with an interactive polygon. The user then moves forward through the frames of a time-lapse movie and manipulates the polygon to define changing boundaries of the same cell. Once cells are segmented and puncta are localized, data for a single cell time-course are automatically combined and stored and the number of puncta identified per cell is displayed in a graph in the lower right-hand corner of the dNEMO GUI. A detailed view of identified puncta can be shown using a dedicated ‘spot inspector tool’, and spots erroneously identified by the wavelet transform (larger objects, vesicular artifacts) can be manually excluded with an interactive removal tool. Results can be saved and reloaded within the dNEMO interface for further analysis. All per-cell and per-spot trajectories are stored in the resulting output files in MATLAB and Excel-compatible formats.

### The *à trous* wavelet transform

Implementation of the *à trous* wavelet algorithm is largely adapted from Izeddin and colleagues (22). Briefly, the raw image is convolved with a matrix to create a wavelet map of the initial image. The L1 kernel is initially populated with values supplied by the third order B-spline (B3) (32). As the level of the transform increases (L2, L3, etc.) zeros are inserted between each of the initial matrix values at increasing amounts. This effectively adds “holes” in the convolution matrix with more zeros inserted as the level of the wavelet transform increases (see Supplemental Figure 1). Subtracting the convolved image from the initial image produces a wavelet map. To identify puncta, a user-defined value is multiplied by the standard deviation of the distribution of pixel intensities in the wavelet map to define a threshold. The threshold is used to classify pixels as either background or foreground. Sub-threshold pixel values are set to 0 and considered background, foreground pixels are associated with puncta and analyzed further in the following watershed segmentation and analysis steps.

### Watershed segmentation

To determine centroids within each identified region within the wavelet map we used the MATLAB (Mathworks) watershed function. Pixel values for the selected wavelet map (L1, L2, etc.) are inverted so that foreground objects act as “catchment basins” for the watershed transform. For each punctate structure identified by the watershed transform, properties of interest are measured as defined by the user, including centroid location, size, and intensity of the punctum, among others (see guide packaged with software).

### Local background correction

Background correction is performed locally for individual puncta in the source image. A binary mask is created representing the regions identified by the *à trous* wavelet transform and dilated by a user-defined number of pixels (Supplemental Figure 3). Pixel intensity values are collected from the annular ring around each punctum and used as the punctum’s local background. An additional user-defined parameter can be assigned to offset the inner diameter of the annular ring. The buffer region excludes the background pixels that immediately surround the punctum and may contain out-of-focus light from the fluorescent source. The background pixel values measured in the annular ring for a punctum are averaged and subtracted from each pixel identified within the punctum in the source image.

### Cell segmentation

Segmentation of individual cells is performed by the user using an interactive polygon tool. This polygon can be further adjusted by the user in subsequent frames to account for morphology changes and cell movements over time. Cell segmentation uses the keyframing approach described above.

### Simulated images with diffraction-limited objects

Three-dimensional matrices of size 512 × 512 × 64 (X, Y, Z) were populated with zeros. A polygon containing pixel values comparable with intensity values found in cellular regions in fluorescence microscopy images was inserted into every z-plane of each matrix to approximate slow-varying background fluorescence, such as unbound dyes or fluorescent proteins that do not reside in puncta. Pixel values within each polygon were increased moving from the polygon’s edges towards the polygon’s center and eventually plateaued at some constant value halfway between the polygon’s edges and centroid. A small Gaussian smoothing filter was applied to each pixel associated with the inserted polygon. Theoretical point spread functions (PSFs) produced by the software package PSFGenerator (33) were individually inserted into the three-dimensional matrix within the bounds of the previously created polygon. To emulate punctate structures whose fluorescence would not be uniform when captured under a microscope, individual PSF pixel values were multiplied by a brightness factor randomly selected in the range [0.5, 1] prior to insertion. Once the matrix contained both simulated cellular regions and theoretical PSFs, multiple instances of the same matrix were produced with increasing noise. Gaussian noise was introduced to the matrix from a distribution with increasing standards of deviation up to 300. The MATLAB code to produce a set of simulated images of some dimensions [height, width, depth] with N number of theoretical PSFs and Gaussian noise along S standards of deviation is provided alongside the dNEMO application here: https://github.com/recleelab.

### Comparison with FISH-quant results

Simulated images containing theoretical point spread functions (PSFs) approximating diffraction-limited objects of known coordinates and intensities were analyzed separately using dNEMO and FISH-quant (27). Puncta were identified in both dNEMO and FISH-quant using the tools available in the “Spot Filter” or “Spot Detection” components, respectively. Additional post-processing of the puncta identified using FISH-quant was performed using the “Thresholding” tool supplied in FISH-quant. Puncta were considered successfully identified if the measured centroid was within a Euclidean distance of 2 of the true centroids. Error rates for puncta localization accuracy were determined as the simulated image was subjected to Gaussian noise of increasing standards of deviation (Supplemental Figure 4). The error rate was determined as the number of false positives and number of false negatives found in each image over the total number of signals present within the image (either 1500, 2500, or 5000 theoretical PSFs; Supplemental Figure 4a).

### Benchmarking dNEMO results

The same fixed-cell smFISH 3D image (1024 × 1024 × 45) image was analyzed with dNEMO and FISH-quant on a 2015 MacBook Pro laptop (16 GB RAM, 2.5 GHz processor) or a Intel® Xeon® PC (128 GB RAM, 2.3 GHz processor). Times were measured using native methods in MATLAB for the core punctum detection algorithms and fluorescence intensity quantification. Reported times do not include image pre-processing steps in FISH-quant (background correction and background region assignment, among others), image loading times, or other user-GUI interactions. The smFISH image was analyzed 5 times in each application for statistical comparisons (Supplemental Figure 4c).

### Comparing smFISH transcripts identified in raw and deconvolved images

NFKBIA transcripts detected by smFISH in HeLa cells were obtained from a previous study (25) and were deconvolved with SoftWoRx using hardware specifications for the DeltaVision microscope (Applied Precision, GE Healthcare Life Science) used for the original image acquisition. Both images were analyzed with dNEMO and mean intensities for the same puncta were compared between images. We show both the uncorrected mean intensity and mean intensity corrected for local background (Supplemental Figure 4e). The R^2^ value for identified puncta is improved significantly (Supplemental Figure 4f), demonstrating the accuracy for measurement of background-corrected puncta.

## SUPPLEMENTAL INFORMATION

Supplemental information includes four figures and two movies can be found with this article online.

## AUTHOR CONTRIBUTIONS

Conceptualization, R.E.C.L.; Methodology, R.E.C.L., G.J.K., and N.S.; Software G.J.K., N.S., and R.E.C.L.; Image Acquisition, J.A.C., Y.G., Q.Z., and R.E.C.L.; Software Testing, J.A.C., Y.G., Q.Z.; Writing – Original Draft, G.J.K. and R.E.C.L.; Writing – Review & Editing, R.E.C.L., G.J.K., J.A.C., Y.G., and N.S.; Visualization, R.E.C.L. and G.J.K; Funding Acquisition, R.E.C.L.; Supervision, R.E.C.L.

## ACKNOWLEDGMENTS

We thank Sanjana Gupta, Chaitanya Mokashi, and David Schipper for helpful discussions. We also thank Suzanne Gaudet for the use of smFISH images. This work was funded by NIH grant (R35-GM119462) to R.E.C.L.

