## Supplemental Information for "dNEMO: a tool for quantification of mRNA and punctate structures in time-lapse images of single cells"

**a**

$$\begin{bmatrix} \frac{1}{256} & \frac{1}{64} & \frac{3}{128} & \frac{1}{64} & \frac{1}{256} \\ \frac{1}{64} & \frac{1}{16} & \frac{3}{32} & \frac{1}{16} & \frac{1}{64} \\ \frac{3}{128} & \frac{3}{32} & \frac{9}{64} & \frac{3}{32} & \frac{3}{128} \\ \frac{1}{64} & \frac{1}{16} & \frac{3}{32} & \frac{1}{16} & \frac{1}{64} \\ \frac{1}{256} & \frac{1}{64} & \frac{3}{128} & \frac{1}{64} & \frac{1}{256} \end{bmatrix}$$

**b**

$$\begin{bmatrix} \frac{1}{256} & 0 & \frac{1}{64} & 0 & \frac{3}{128} & 0 & \frac{1}{64} & 0 & \frac{1}{256} \\ 0 & 0 & 0 & 0 & 0 & 0 & 0 & 0 & 0 \\ \frac{1}{64} & 0 & \frac{1}{16} & 0 & \frac{3}{32} & 0 & \frac{1}{16} & 0 & \frac{1}{64} \\ 0 & 0 & 0 & 0 & 0 & 0 & 0 & 0 & 0 \\ \frac{3}{128} & 0 & \frac{3}{32} & 0 & \frac{9}{64} & 0 & \frac{3}{32} & 0 & \frac{3}{128} \\ 0 & 0 & 0 & 0 & 0 & 0 & 0 & 0 & 0 \\ \frac{1}{64} & 0 & \frac{1}{16} & 0 & \frac{3}{32} & 0 & \frac{1}{16} & 0 & \frac{1}{64} \\ 0 & 0 & 0 & 0 & 0 & 0 & 0 & 0 & 0 \\ \frac{1}{256} & 0 & \frac{1}{64} & 0 & \frac{3}{128} & 0 & \frac{1}{64} & 0 & \frac{1}{256} \end{bmatrix}$$

**c**

$$\begin{bmatrix} \frac{1}{256} & 0 & 0 & 0 & \frac{1}{64} & 0 & 0 & 0 & \frac{3}{128} & 0 & 0 & 0 & \frac{1}{64} & 0 & 0 & 0 & \frac{1}{256} \\ 0 & 0 & 0 & 0 & 0 & 0 & 0 & 0 & 0 & 0 & 0 & 0 & 0 & 0 & 0 & 0 & 0 \\ 0 & 0 & 0 & 0 & 0 & 0 & 0 & 0 & 0 & 0 & 0 & 0 & 0 & 0 & 0 & 0 & 0 \\ 0 & 0 & 0 & 0 & 0 & 0 & 0 & 0 & 0 & 0 & 0 & 0 & 0 & 0 & 0 & 0 & 0 \\ \frac{1}{64} & 0 & 0 & 0 & \frac{1}{16} & 0 & 0 & 0 & \frac{3}{32} & 0 & 0 & 0 & \frac{1}{16} & 0 & 0 & 0 & \frac{1}{64} \\ 0 & 0 & 0 & 0 & 0 & 0 & 0 & 0 & 0 & 0 & 0 & 0 & 0 & 0 & 0 & 0 & 0 \\ 0 & 0 & 0 & 0 & 0 & 0 & 0 & 0 & 0 & 0 & 0 & 0 & 0 & 0 & 0 & 0 & 0 \\ 0 & 0 & 0 & 0 & 0 & 0 & 0 & 0 & 0 & 0 & 0 & 0 & 0 & 0 & 0 & 0 & 0 \\ \frac{3}{128} & 0 & 0 & 0 & \frac{3}{32} & 0 & 0 & 0 & \frac{9}{64} & 0 & 0 & 0 & \frac{3}{32} & 0 & 0 & 0 & \frac{3}{128} \\ 0 & 0 & 0 & 0 & 0 & 0 & 0 & 0 & 0 & 0 & 0 & 0 & 0 & 0 & 0 & 0 & 0 \\ 0 & 0 & 0 & 0 & 0 & 0 & 0 & 0 & 0 & 0 & 0 & 0 & 0 & 0 & 0 & 0 & 0 \\ 0 & 0 & 0 & 0 & 0 & 0 & 0 & 0 & 0 & 0 & 0 & 0 & 0 & 0 & 0 & 0 & 0 \\ \frac{1}{64} & 0 & 0 & 0 & \frac{1}{16} & 0 & 0 & 0 & \frac{3}{32} & 0 & 0 & 0 & \frac{1}{16} & 0 & 0 & 0 & \frac{1}{64} \\ 0 & 0 & 0 & 0 & 0 & 0 & 0 & 0 & 0 & 0 & 0 & 0 & 0 & 0 & 0 & 0 & 0 \\ 0 & 0 & 0 & 0 & 0 & 0 & 0 & 0 & 0 & 0 & 0 & 0 & 0 & 0 & 0 & 0 & 0 \\ 0 & 0 & 0 & 0 & 0 & 0 & 0 & 0 & 0 & 0 & 0 & 0 & 0 & 0 & 0 & 0 & 0 \\ \frac{1}{256} & 0 & 0 & 0 & \frac{1}{64} & 0 & 0 & 0 & \frac{3}{128} & 0 & 0 & 0 & \frac{1}{64} & 0 & 0 & 0 & \frac{1}{256} \end{bmatrix}$$

**Figure S1. Wavelet convolution kernels used to enhance diffraction limited and near diffraction-limited puncta from microscopy images.** Kernel matrices used for **(a)** L1, **(b)** L2, and **(c)** L3 wavelet decomposition. The *a trous* wavelet transform adds progressively more zeroes, referred to as ‘holes’, at each level.

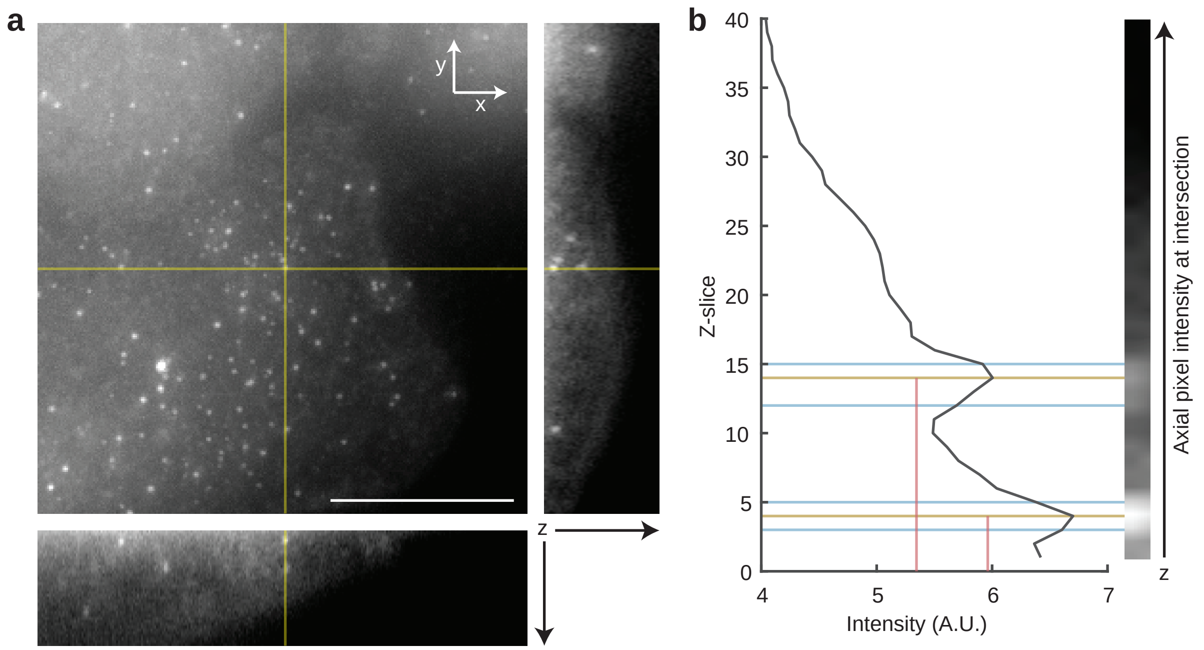


**Figure S2. Puncta with axial overlap are individually resolved. (a)** Maximum intensity projection of a representative *NFKBIA* smFISH fluorescent z-stack from HeLa cells. Axial projections across the indicated yellow lines for x-z (right) and y-z (bottom) planes show two puncta that overlap in x-y but are axially separated. Scale bar is 10 µm. **(b)** Axial pixel intensities at the x-y intersection of the yellow lines in panel ‘a’ (right). The pixel values are measured at each slice of the z-stack (left). Tan lines indicate the axial slice of the centroid for each punctum and teal lines indicate the contiguous z-slices in which the same punctum is observed. Red vertical lines indicate the local background intensity value determined at the axial centroid and used to correct intensity values for each punctum within the 3D image.

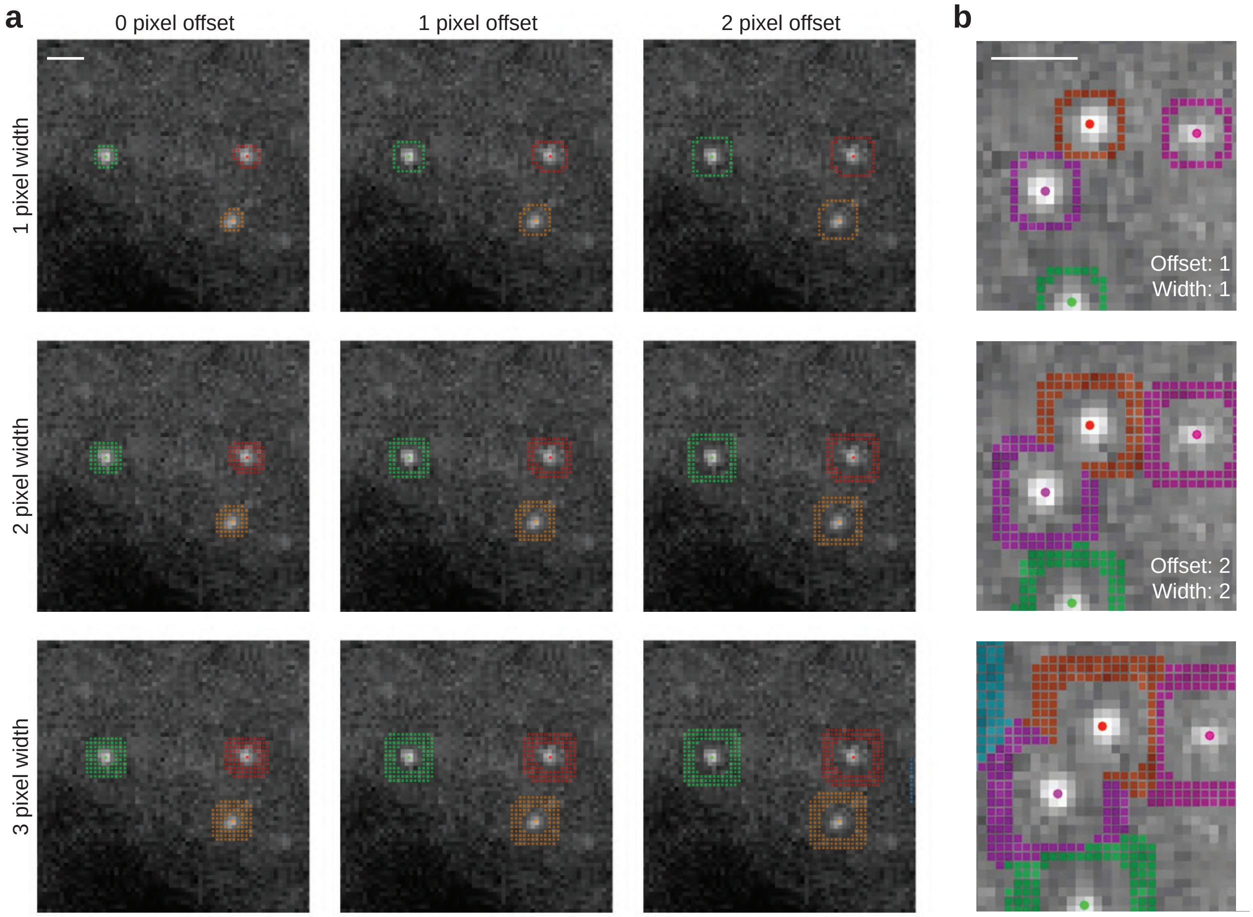


**Figure S3. Local background information is used to correct intensity values measured for each punctum. (a)** For each punctum, pixel intensity values are collected from an annulus with user defined inner radius (offset) and width, defined in pixel units. The mean intensity value of pixels bound by the annulus is subtracted from every pixel within the associated punctum in the source image to correct for the local background. Because each punctum is measured relative to local background, images do not require pre-processing before analysis with dNemo. **(b)** Pixels in the vicinity of a spot will be excluded from annuli of all nearby spots. Similarly, when background annuli overlap, background pixels will be associated with the nearest spot only, as determined by its centroid. Images of smFISH against the *NFKBIA* transcripts in HeLa cells. Scale bars equal 1 µm.

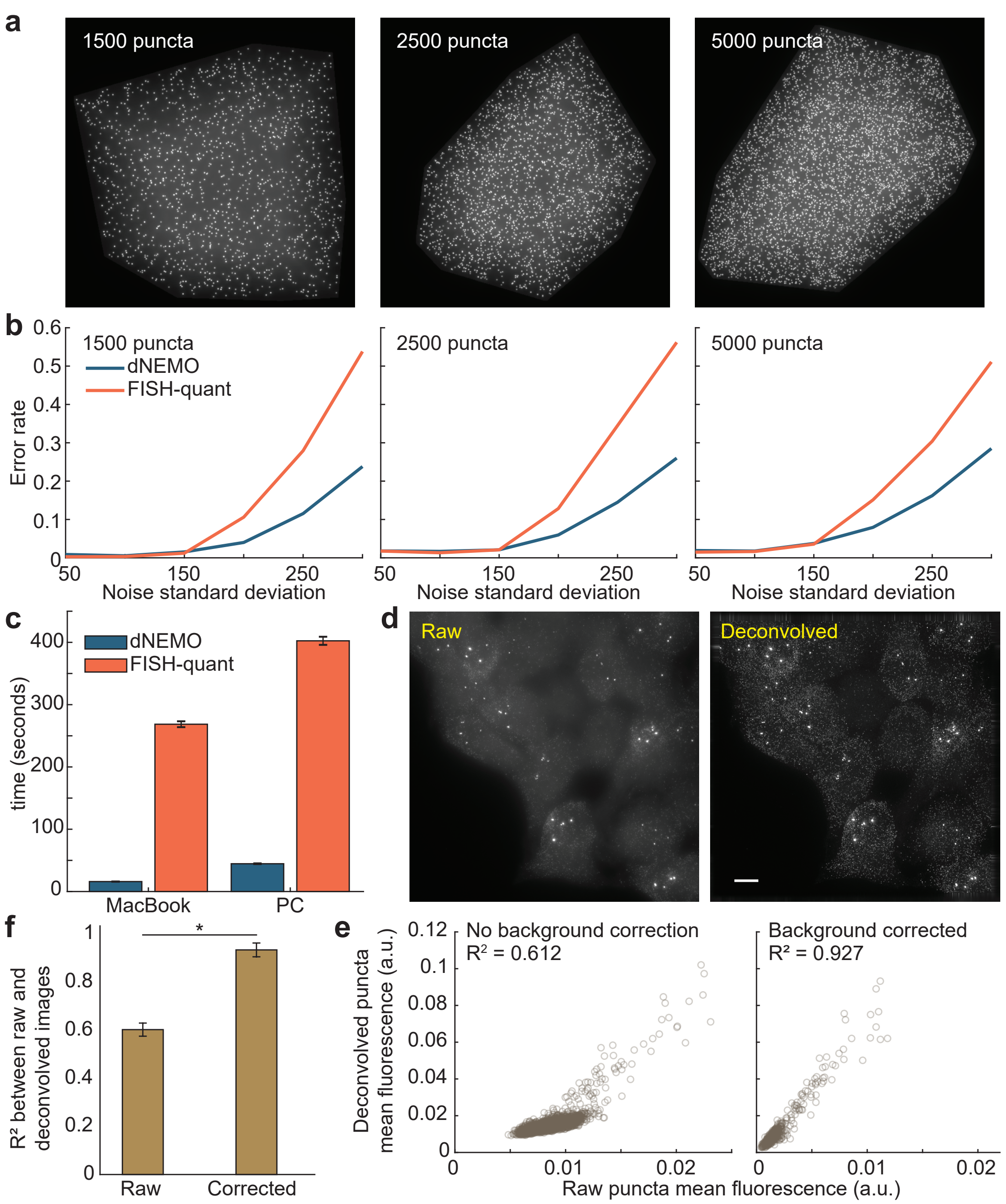


**Figure S4. Benchmarking results for dNEMO.** (a) Maximum intensity projection (MIP) for simulated 3D images using a theoretical point spread function for the Deltavision microscope over a slow-varying background with Gaussian noise. Inset indicates the number of simulated puncta in each image. **(b)** Images were processed using dNEMO and FISH-quant with increasing Gaussian noise. The error rate is the number of false positives and false negatives divided by the total number of puncta in the simulated data. A punctum is accurately detected if the centroid falls within 2 pixels of the true value in simulated data. **(c)** Bar graph for single-core runtime comparison between a MacBook and PC with comparable single-core speeds (2.5 versus 2.3 GHz respectively). The same 3D smFISH image was analysed by dNEMO and FISH-quant as indicated (n = 5). **(d)** MIP for smFISH of TNF-induced *NFKBIA* gene expression. The raw image (left) and deconvolved image (right) using SoftWoRx are shown. Scale bar 10uM. **(e)** Punctum intensity values for both images in panel (d) were compared, calculated without (left) and with dNEMO background correction (right). $R^2$ values between raw and deconvolved images improve significantly ($p < 0.0002$, t test) with background correction, quantified in **(f)** for 3 representative images. Error bars ± S.D. for all.
